# Solution-phase indexing by kinetic confinement enables rapid, simple, and instrument-free single cell transcriptional profiling

**DOI:** 10.1101/2024.11.01.621570

**Authors:** P. Marafini, DG. Smith, AR. Lamstaes, RE. Contreras, I. Williams, I. West, O. Ambridge, V. Sanders-Brown, E. Intaite, CY. Hii, BCC. Hume, U. Munagala, W. Plumbly, FL. Brown, Y. Shlyakhtina, L. Woods, JA. Bibby, L. Williams, JHM. Yang, B. Steffy, L. Zawada, JW. Harger, D. McKenzie, AG. Laing, MJT. Stubbington, LB. Edelman

## Abstract

Existing tools for single cell genomics require complex physical frameworks for the indexing of cellular nucleic acids, including proprietary instrumentation, droplet emulsions, and laborious combinatorial indexing schemes. The complexity and cost of these tools significantly constrains the use of single cell technologies across basic and translational research.

Here, we describe an instrument-free method that uses novel, bifunctional indexing reagents to deliver index sequences directly to single cells followed by a biophysical process known as ‘Kinetic Confinement’ to perform high-fidelity indexing of target molecules across thousands of single cells simultaneously in single-tube, solution-phase reactions.

Kinetic Confinement enables simple, fast, and flexible single cell experiments, and allows straightforward scaling to very large sample numbers. We anticipate that assays based on Kinetic Confinement will significantly expand the scope, use, and impact of single cell analysis across fundamental and applied research, as well as within therapeutic development and ultimately applied clinical diagnostics.

## Introduction

Single cell RNA sequencing (scRNAseq) was named Method of the Year in 2013^1^ and has, in the intervening years, enabled many highly-impactful discoveries across basic^2^ and translational^3^ research.

Existing methods for scRNAseq all rely on the introduction of a physical, impermeable partition between individual cells during indexing of mRNA molecules from those cells. These physical partitions can include the outer shell of an aqueous microdroplet generated by microfluidics^4–6^ or by bulk emulsification when cells are combined with beads within those droplets^7^; the wall of an individual well within a microwell plate in assays that use split–pool combinatorial indexing approaches^8,9^; or the walls of a nanowell array where cells are deposited by settling^10^.

These partition-based methods are suited to the analysis of large numbers of *cells* from relatively small numbers of individual samples. In addition, the value of single cell genomics has been demonstrated in applications that would, typically, depend on large numbers of *samples* within complex, multifactorial designs such as compound screening within a heterogeneous system of primary cells^11^ or population-scale studies of inter-individual transcriptional variation^12^. However, experiments such as these are not commonplace due to the requirements of existing single cell methods for expensive instrumentation and/or complex workflows that use multi-step liquid handling methods that are unfamiliar even to experienced molecular biologists. This means that a large suite of applications continue to use bulk genomics or orthogonal non-genomics approaches and are missing out on the high-resolution, data-rich insights provided by single cell assays.

Here, we present an approach to scRNAseq where single cell indexing is performed in-solution and without physical partitioning. Instead, single cell indexing is enforced by a combination of a cell-binding indexing reagent along with a viscous lysis and indexing buffer that prevents diffusion and inter-cell crosstalk of mRNA molecules and indexing oligonucleotides. This “Kinetic Confinement” method enables scRNAseq to be performed in standard laboratory plasticware without the need for any specialised instrumentation, nor arduous workflows. Individual samples are never split and recombined, and liquid handling steps consist solely of standard pipetting and magnetic bead cleanups.

scRNAseq by Kinetic Confinement can be performed at low cost on a single sample without the complexity of split–pool methods. In addition, the assay’s simplicity also allows it to scale easily to parallelised experimental designs with tens, hundreds or even thousands of individual samples. Although no instrument is *required* for the assay, its workflow is uniquely amenable to use with automated or semi-automated liquid handling systems should these be necessary to enable ultra-large scale experiments with very many different samples.

We believe that this technology has the potential to transform applications such as pharmaceutical compound screening and to enable experiments that have, as yet, not been possible. Furthermore, rapid, scalable, and automatable data generation methods that provide rich, high-resolution data will be necessary in studies where active learning and artificial intelligence is used to iteratively design, perform, and analyse experiments^13^. This has the potential to accelerate drug discovery campaigns and to drive extensive insights within fundamental research.

## Results

### Kinetic Confinement enables single cell transcriptomic analysis

Single cell RNAseq using Kinetic Confinement (Fig 1a) relies upon two key reagents. The first is a cell pairing and indexing reagent (“CPair”) that comprises a paramagnetic bead of comparable diameter to the majority of mammalian cells. CPair beads are functionalized with both bead-specific Indexing Oligonucleotides (IOs), and with ‘cell-binding molecules’ that allow the beads to bind strongly and non-selectively to cells.

**Figure 1.**
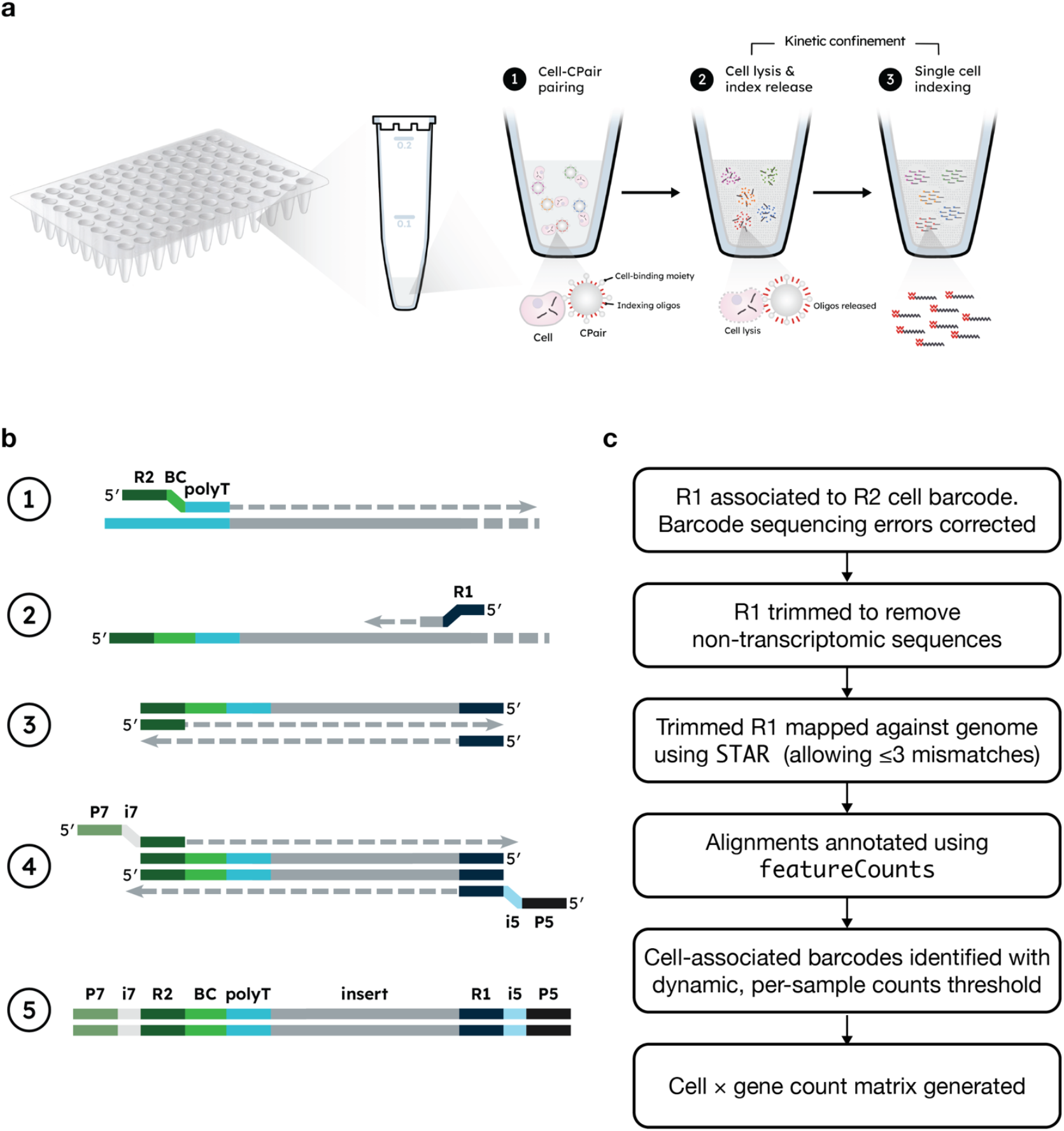
Overview of the SimpleCell 3′ Single Cell Gene Expression Assay a, Single cell indexing is achieved by pairing cells with CPair and then performing lysis and indexing within viscous Kinetic Confinement Bu?er. **b**, Library preparation steps including (1) reverse transcription, (2) second strand synthesis primed at random locations so that each molecule has a unique Second Strand Synthesis Start Site (5S) coordinate, (3) PCR amplification, (4) Second PCR amplification to introduce sequencing adapters and indices, (5) final library ready for sequencing using short-read NGS technologies. **c**, Sequencing data are processed with a custom, open-source computational pipeline that generates a vector of gene expression values for each cell-associated barcode.

Each Indexing Oligonucleotide consists of: i) a poly(dT) sequence for priming of reverse transcription from the polyA tail of messenger RNA molecules; ii) a 26 base-pair barcode drawn from a pool of ∼50,000 unique sequences for indexing cDNA products from each individual cell; iii) A primer annealing site for subsequent PCR amplification steps. All of the Indexing Oligonucleotides for each individual CPair bead are identical, but the barcode sequences differ between CPair beads.

The second key reagent is a viscous “Kinetic Confinement Buffer” (KCB). This buffer restricts diffusion of nucleic acids and also contains a heat-activatable cell lysis agent.

To perform single cell RNAseq, prepared cells are mixed with the CPair reagent such that 1:1 cell–bead binding events are strongly favoured through a combination of optimized cell:bead input ratios, buffer volumes, reaction vessel size and shape, and handling conditions. Strongly bound cell–bead pairs form driven by the cell-binding molecules on the surface of each bead.

Once cells are paired with CPair beads, they are transferred into KCB and subjected to a short high-temperature incubation followed by slow cooling in a standard thermocycler. The high temperature causes cell lysis and also releases indexing oligonucleotides from each CPair bead. The ramped cooling allows indexing oligonucleotides to hybridize to mRNA molecules released from lysed cells through their polydT sequences.

Single cell behaviour is maintained during the indexing process by the high viscosity of the Kinetic Confinement Buffer. This constrains diffusion of both mRNA molecules and indexing oligonucleotides, such that 1) the nucleic acids from a single cell–bead pair exist in a high local concentration, which drives high-efficiency indexing of cellular nucleic acids from within that pair, and 2) indexing oligonucleotides and mRNA from each given cell–bead pair remain physically distant from other cell–bead pairs, and thus mRNA is significantly more likely to be indexed by an oligonucleotide released from the CPair bead to which that cell was bound.

Kinetic Confinement favours the kinetics of intra-pair cell indexing, and suppresses the kinetics of inter-pair indexing, thus enabling high-fidelity single-cell indexing in single-tube, solution-phase reactions.

Single cell indexing of mRNA molecules is followed by reverse transcription, second strand synthesis and library preparation steps to generate libraries of DNA molecules that can be sequenced using short-read NGS technologies (Fig 1b).

Post-reverse transcription second strand synthesis is performed using primers that anneal to the first cDNA strand at diverse, random locations. This results in each second strand synthesis product having a unique Second Strand Synthesis Start Site (“5S”). Our custom bioinformatic pipeline (Fig 1c) uses the genomic mapping coordinates of each 5S to identify and collapse sequencing reads that arise from PCR duplication events downstream of initial mRNA indexing. This allows accurate expression quantification without using unique molecular identifiers (Supplementary Fig 1).

Kinetic Confinement enables rapid processing of single cell suspensions to sequencer-ready libraries with the workflow for eight samples taking 8h of total time, of which 4h10m is hands-on.

### Kinetic Confinement performance in species-mixing experiments

We assessed the performance of Kinetic Confinement-based single cell RNAseq by analysing a mixture of 2500 mouse (NIH3T3) and 2500 human (HeLa) cells as input to the assay followed by sequencing the library to an average depth of 39,886 reads/cell. We used our computational pipeline (Fig 1c) to identify the CPair barcode associated with each read and to map transcriptome reads to both the mouse and human genomes using splice-aware mapping. Reads that mapped to transcripts were deduplicated using their 5S sites and these counts were used to estimate gene expression for each gene associated with each CPair barcode.

96.5% of reads contained valid CS Genetics cellular barcodes and, of these, 41% mapped confidently to the human transcriptome while 44% mapped confidently to the mouse transcriptome. We used the per-barcode distribution of 5S counts for human and mouse genes to identify barcodes arising from cell–CPair pairs (Fig 2a) and to classify these barcodes as human, mouse, or mixed species (Fig 2b). We recovered 4376 cell-associated barcodes, of which 1926 (44%) were human, 2097 (48%) were mouse, and 353 (8%) were mixed. This indicates that kinetic confinement maintains single cell indexing behaviour without the need for physical partitioning of cells.

**Figure 2.**
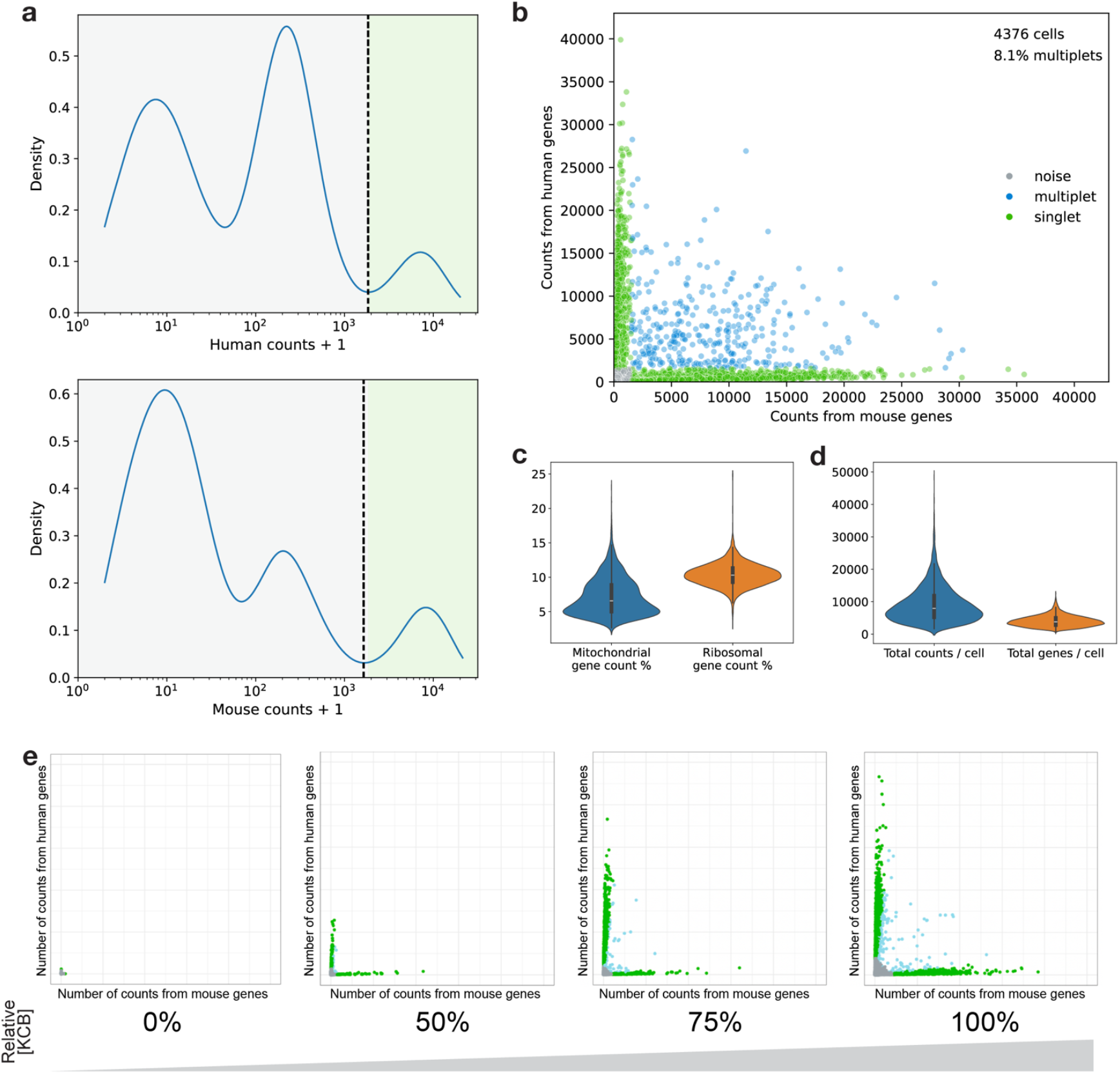
Species-mixing experiments to demonstrate single cell RNA-sequencing using Kinetic Confinement. **a**, Identification of cell-associated barcodes. The 5S count distribution for majority-human (top) or majority-mouse (bottom) cells is used to dynamically determine a threshold(dotted line) to separate cell-associated CPair barcodes (green area) from those that are not (grey area). **b**, Assessment of multiplet rate using mixed species input. For each CPair barcode, the number of counts associated with the mouse genome and the human genome are determined and the cell-caller thresholds are used to classify each barcode as noise (grey points), a single cell (green points), or a mixed-species multiplet (blue points). **c**, Distributions of per-cell quality control metrics: percentage of reads derived from the mitochondrial genome (blue), and percentage of reads derived from genes encoding ribosomal proteins (orange). **d**, Distributions of the number of per-cell 5S counts (blue) and genes detected (orange). **e**, Effects of increasing KCB concentration upon single cell indexing. A species-mixing experiment was performed with the lysis and indexing step in the usual KCB formulation (far right) or with increasing dilutions of KCB in PBS.

Within the cells, a median of 6.6% of reads were assigned to genes carried on the mitochondrial genome while 10.3% derived from genes encoding ribosomal proteins (Fig 2c; Supplementary Figure 2)

We detected a median of 3810 genes per cell-associated barcode (Fig 2d; Supplementary Figure 2). These values were derived from a median of 7967 5S counts per cell-associated barcode.

We confirmed the importance of Kinetic Confinement Buffer for single cell analysis by performing species-mixing experiments using diluted KCB formulations from 0% KCB (solely PBS) up to the full KCB concentration (Fig 2e). Without KCB we observed essentially no cellular signal and this increases up to the expected single cell behaviour with 100% KCB.

### Kinetic Confinement enables fixation-free storage of single cell samples prior to analysis

Kinetic Confinement Buffer has a cryoprotective effect upon frozen cells which enables the workflow to be paused after just two hours. This is useful in cases where cell samples are ready later in the day so that library preparation can be safely completed on the following day. Furthermore, the use of single cell sequencing technologies can be impeded by the need to collect and process multiple cell samples all at the same time. This would become particularly challenging with the high number of samples per experiment that is compatible with Kinetic Confinement assays.

To confirm whether the cryoprotection of KCB would allow longer-term storage of paused samples, we investigated the stability of paired cell–CPair samples stored at -80C in KCB. We generated mixed-species cell samples that were prepared on the same day and processed through the assay until the transfer of paired cells into KCB. These samples were then frozen and stored at -80C overnight (“Day 0”) or for up to four weeks prior to performing the remaining library preparation and sequencing steps (Fig 3a). Libraries were sequenced to a median depth of 4,000 reads/cell.

**Figure 3.**
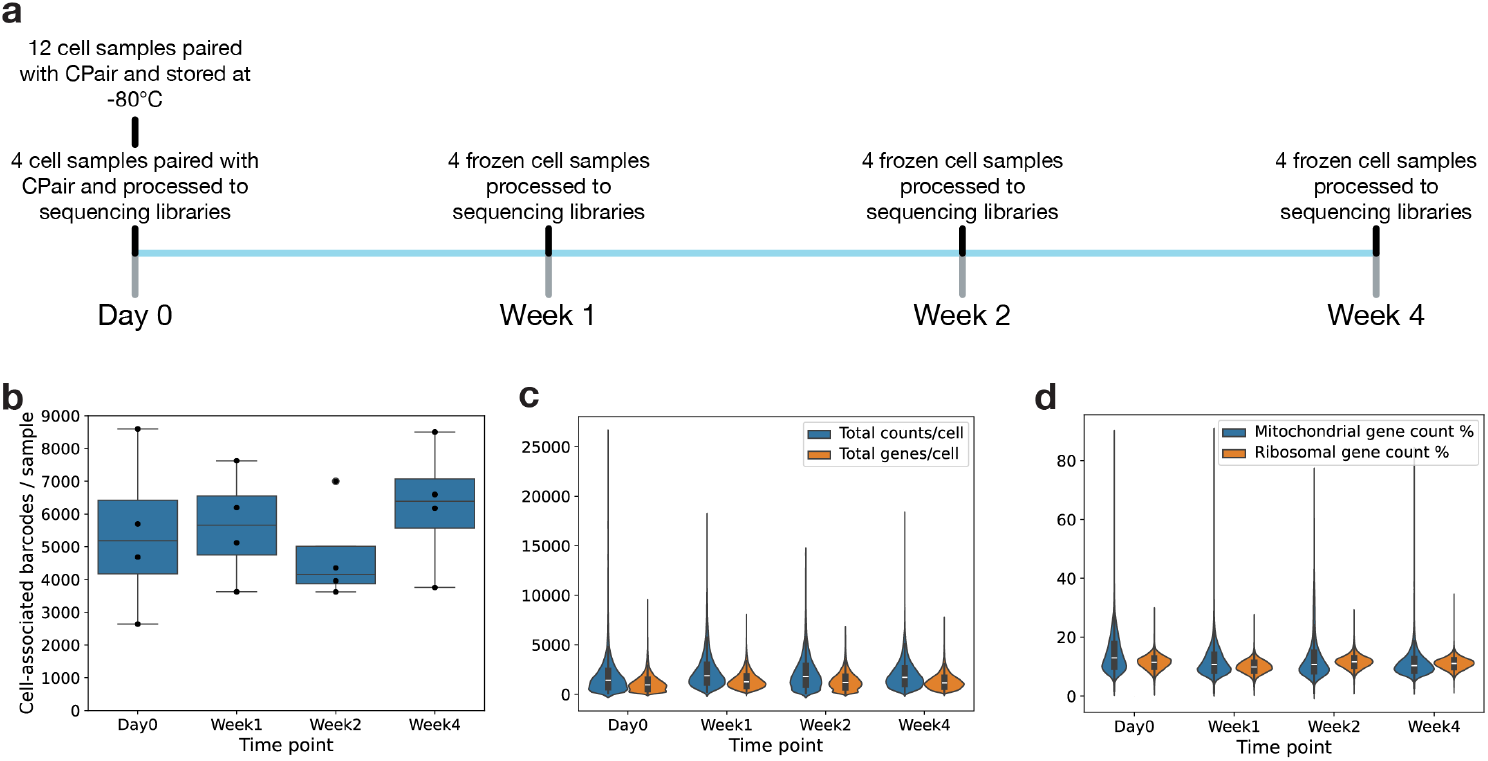
Stability of cells paired with CPair frozen in KCB. **a**, Experimental setup. 16 replicate mixed-species cell-line samples were paired with CPair and transferred to KCB on Day 0. Four samples were processed to single cell RNA-sequencing libraries immediately while 12 others were stored at -80°C for up to four weeks with four frozen samples being processed to sequencing libraries each week. **b**, Number of cell-associated barcodes detected in each sample at each timepoint. **c**, Distributions of the number of per-cell 5S counts (blue) and genes detected (orange) at each timepoint. **d**, Distributions of per-cell quality control metrics at each timepoint: percentage of reads derived from the mitochondrial genome (blue), and percentage of reads derived from genes encoding ribosomal proteins (orange).

The Day 0, Week 1, Week 2, and Week 4 samples demonstrated little storage time-dependent difference in cell recovery, per-cell sensitivity, mitochondrial read %, or ribosomal read % (Fig 3b) indicating that storage for four weeks does not lead to degradation in data quality.

### Single cell analysis of heterogeneous primary cell populations

Single cell transcriptomic assays are commonly used to investigate the composition and gene expression of heterogeneous primary cell populations. To confirm the utility of Kinetic Confinement for this application we first assessed its ability to identify immune cell populations within peripheral blood mononuclear cells (PBMCs). We processed four individual samples of 8,500 PBMCs each, purified from a single donor and sequenced them to a median depth of 39,450 reads/cell

After quality filtering, we assigned 15,650 CPair barcodes to cells across all four samples with a median of 3,873 cells per sample. Within these cells, the median number of detected 5S counts was 2,353 per cell and the median detected number of genes was 1,457 per cell (Supplementary Figure 2).

We identified cell types and subtypes within the PBMC population by performing graph-based clustering analysis (Fig 4a). We then identified differentially expressed marker genes that were associated with each cluster and used these along with knowledge of expected PBMC subtype marker genes to annotate cellular identity in each cluster (Fig 4b, Supplementary Fig 3). We identified all the expected major cell types and observed specific subtypes within the T cell and monocyte clusters. The proportions of each cell type within this dataset were within the expected ranges for PBMCs from a healthy donor (Fig 4c).

**Figure 4.**
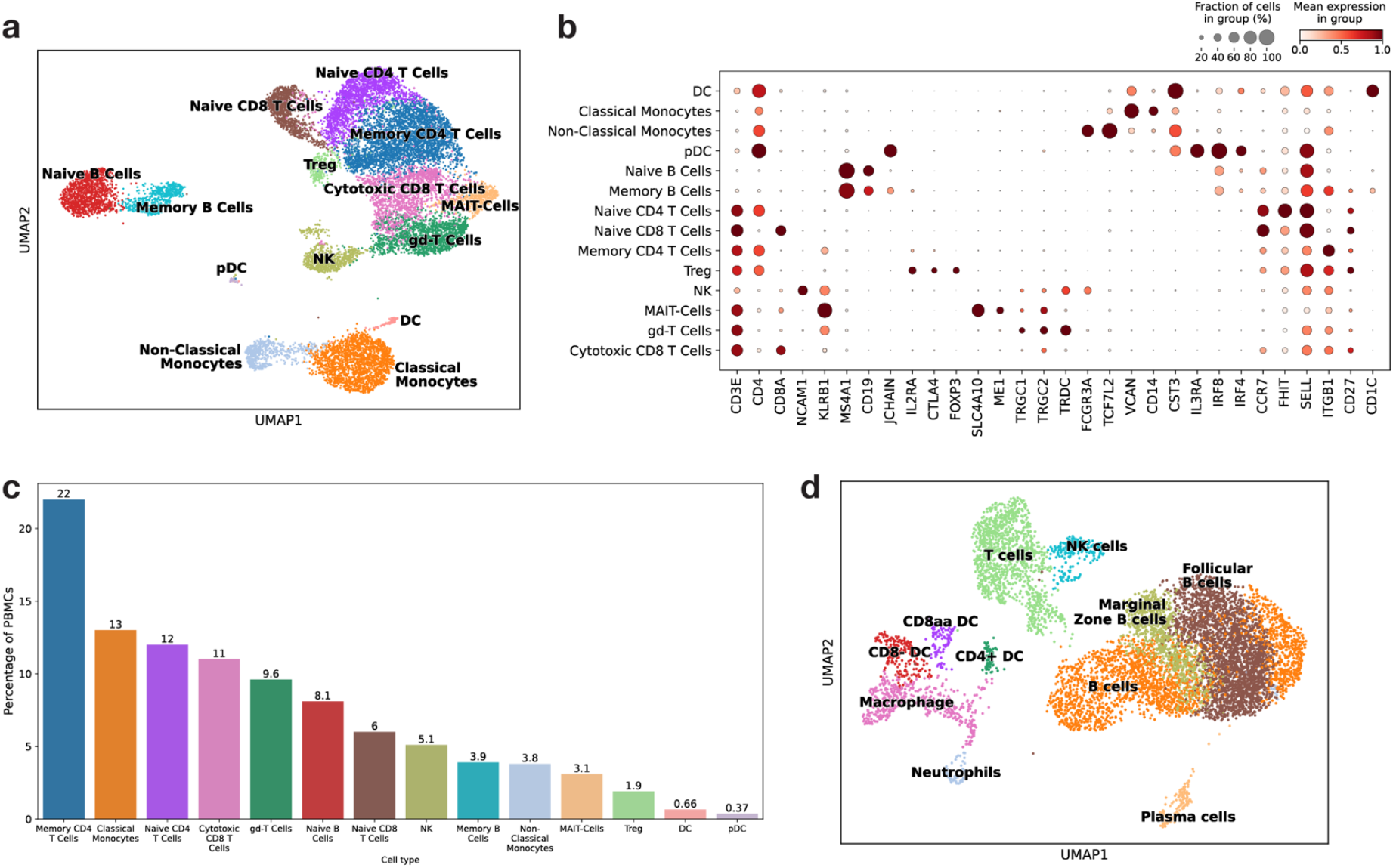
Detection of cell subtypes within heterogeneous mixtures from blood and solid tissue a, Graph-based clustering and cell-type annotation of cells within human PBMC samples.**b**, Expression of a selected set of cell type-specific marker genes within each annotated cluster **c**, Proportions of each cell type identified within the PBMC data **d**, Graph-based clustering and cell-type annotation of cells within dissociated mouse spleen samples

We also used dissociated mouse spleen cells as input into our assay to demonstrate compatibility with disaggregated solid tissue samples. Clustering analysis of these cells again showed recovery of expected cell types at typical proportions (Fig 4d).

### Large-scale analysis of many samples in parallel

It is often not feasible to process large numbers of blood samples to purify PBMCs due to workflow and time constraints. This can prevent the use of single cell assays in population-scale projects that could generate data from thousands of individual donors. In these situations, it can be significantly easier to use whole-blood samples without extensive additional processing. We applied our kinetic confinement approach to whole blood samples that had undergone red blood cell lysis followed by dead cell removal using magnetic beads. To illustrate the scale of sample analysis that is possible with this method, we processed 96 individual frozen whole blood samples in a single day through to the fixation-free pause point whereby cell–CPair mixtures were stored at -80C overnight. On the following day, all 96 samples were processed through to sequenceable libraries by a single operator using a 96-channel pipetting instrument.

To enable us to assess within-assay reproducibility, the 96 individual samples were drawn from 48 individual donors such that each donor blood draw was represented by two samples in the randomized final set.

The single cell RNAseq data demonstrated successful cell capture and library generation from all 96 samples (Fig 5a) with very little technical batch effects (Fig 5b). Gene expression quantification was enabled high-resolution annotation of cell types within the samples (Fig 5C).

**Figure 5.**
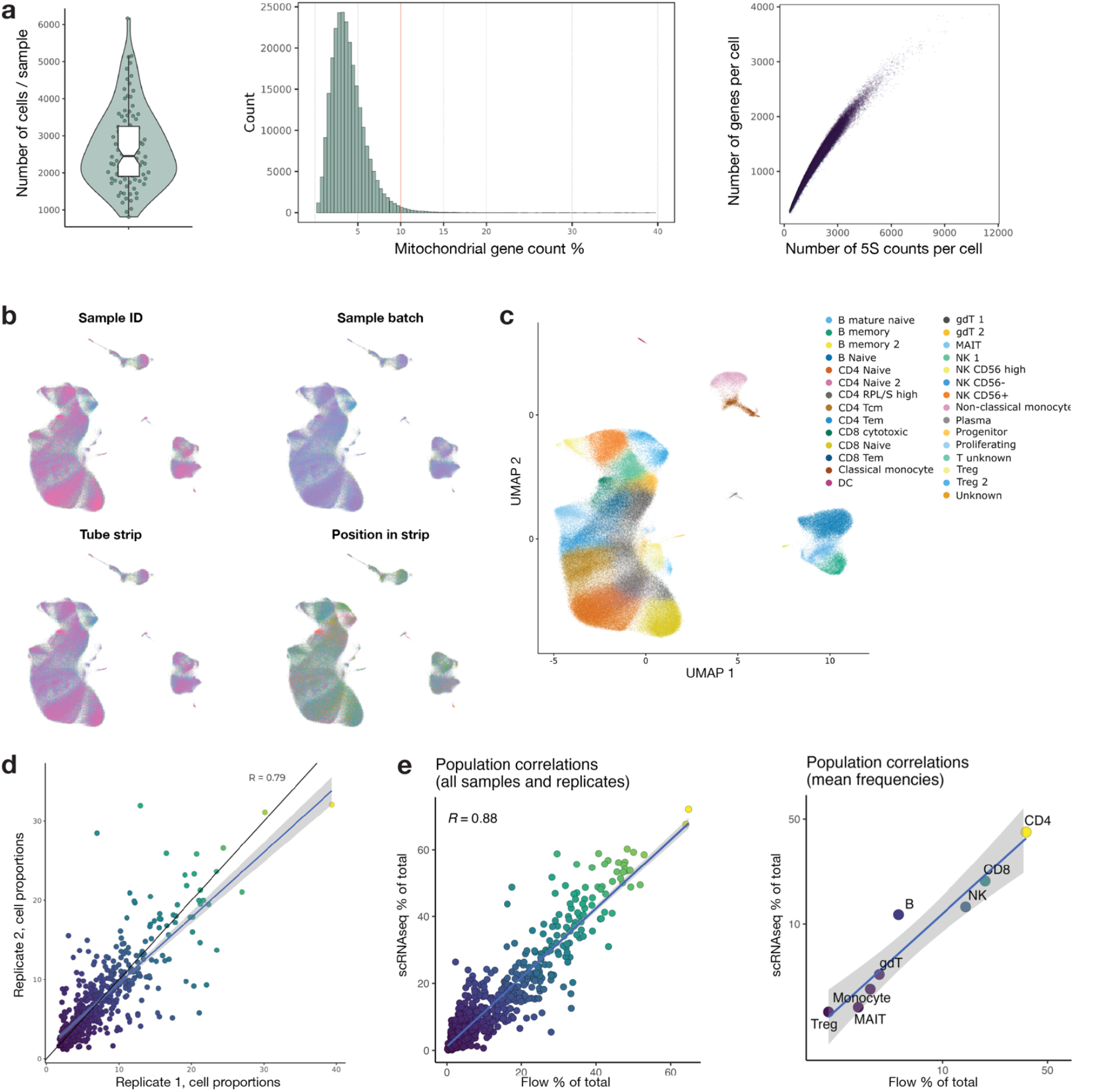
Highly-parallel analysis of 96 individual whole-blood samples. **a**, Quality control metrics showing number of cells detected in each sample (left), distribution of the proportion of genes deriving from the mitochondrial genome in each sample (middle), and the relationship between 5S counts and genes detected within every cell (right). **b**, Dimensionality reduction plots coloured according to possible causes of batch effect within the dataset. **c**. Graph-based clustering and cell-type annotation of cells within all 96 whole-blood samples. **d**, Concordance between cell type population proportions between replicate aliquots from the same original blood donation. **e**, Concordance of identified cell type population proportions between single cell RNA seq and spectral flow cytometry.

We then compared the proportion of each cell type identified within each paired sample from a given replicate and confirmed a high level of between-replicate correlation (Fig 5d).

In addition, each sample was analysed by spectral flow cytometry to provide an orthogonal, ground truth assessment of the cell types present. We investigated whether our kinetic confinement approach was able to capture different cell subtypes in an unbiased manner by comparing the cell type proportions identified in the scRNAseq data with those identified by flow cytometry. There was significant concordance between the proportions identified by these two independent methods (Fig 5e)

### Selective capture of specific cell types from heterogeneous cell mixtures

The cell-binding properties of CPair enable it to be selectively directed towards cell types that express a specific cell surface marker. Here, we replaced the cell-agnostic binding molecule used above with antibodies that bind to one of three different cell surface proteins: CD45 (to target all leukocytes), CD3E (to target T cells), or CD19 (to target B cells). We used three aliquots of a heterogeneous PBMC mixture purified from a single blood sample and performed the Kinetic Confinement assay using each of the three antibody targeting approaches separately. We then performed clustering analysis of all the data combined (Fig 6) and annotated cell types as above.

**Figure 6.**
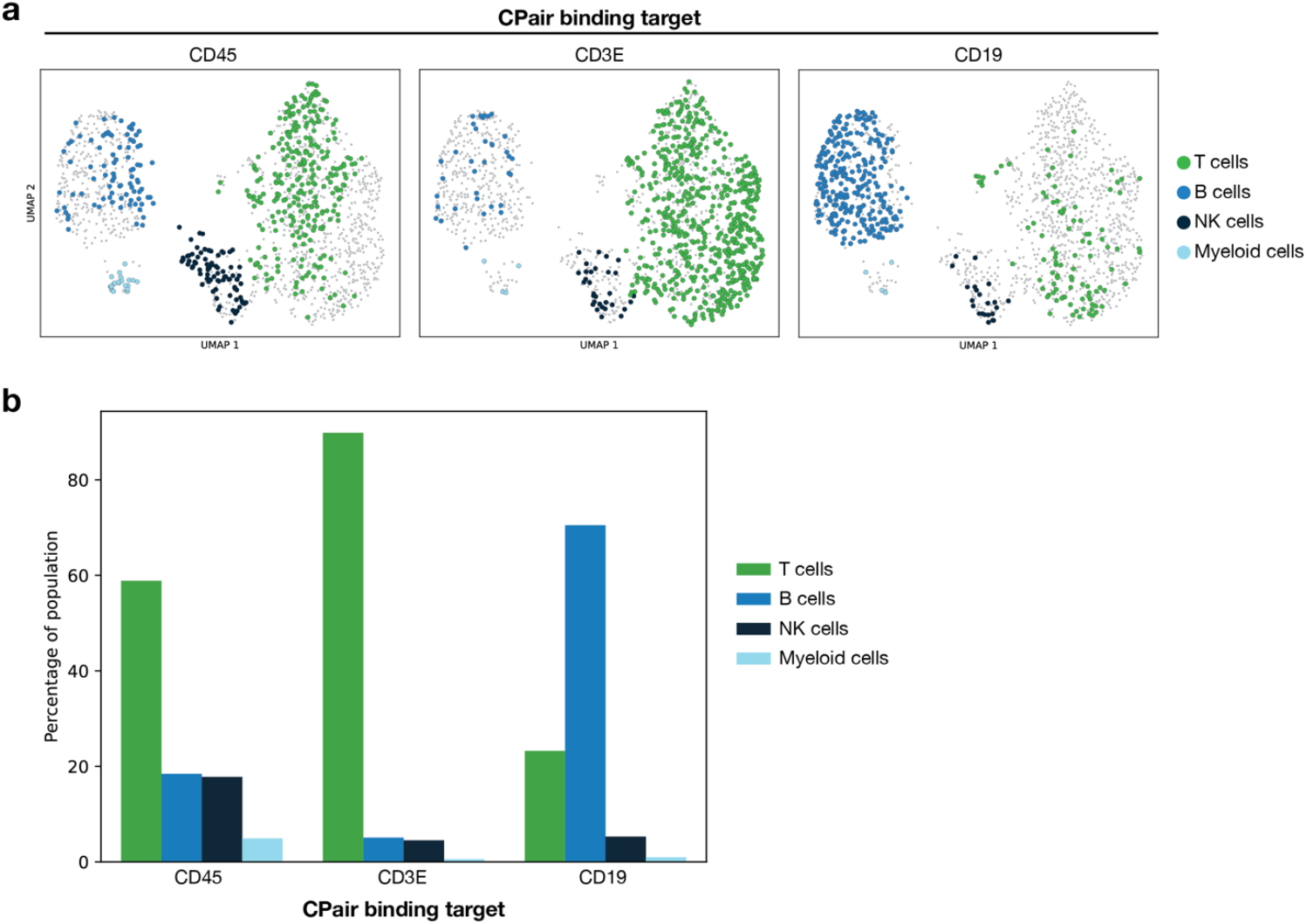
Selective capture of specific cell types with altered CPair binding chemistry. **a**, UMAP dimensionality-reduction and clustering plots from three experiments using separate aliquots of PBMCs from the same donor and where the CPair cell-binding molecule was an antibody targeted against CD45 (left), CD3E (centre), or CD19 (right). All data were analysed together and then plotted separately according to the targeting antibody. Coloured points indicate cells that were present in a given sample whilst grey points indicate cells present in the other two samples. **b**, Proportions of each cell type present within each sample.

Data from CD45-targeted CPair beads showed that all leukocyte populations could be detected in their expected proportions. CD3E-targeted CPair produced enrichment of T cells from 60% to 90% while CD19-targeted reagents gave an enrichment of B cell from 18% to 71%.

## Discussion

Single cell genomics has been a transformative technique for understanding biology for over a decade. As the technology has matured, we have begun to move to a new era where observational “atlassing” studies are being increasingly replaced by experiments that use single cell methods to investigate cell function through various perturbations or due to the impact of population-scale variation. Asking these questions necessitates the use of large numbers of individual samples rather than large numbers of cells from a small number of samples. Existing single cell technologies are not well-matched to these needs due to their need for expensive instrumentation, complex workflows, limits on the number of samples that can be processed in a given experiment, and/or their incompatibility with high-throughput automated liquid handling.

Our approach of Kinetic Confinement replaces partition-based single cell indexing with a solution-phase approach that is uniquely compatible with standard laboratory plasticware and molecular biology techniques. Kinetic Confinement also enables experiments with one to thousands of samples and provides a direct path to use with standard automated NGS liquid handling robots for ultra-large scale experiments.

We have demonstrated the compatibility of single cell indexing by Kinetic Confinement in cell lines and primary cell samples from blood and solid tissue. We showed that Kinetic Confinement can be used to perform scRNAseq from large numbers of whole blood samples without the need for further purification of PBMC populations. In all cases we demonstrated the ability of our approach to identify relevant biological signals within heterogeneous cell mixtures at single cell resolution. We confirmed that it is possible to freeze cells in Kinetic Confinement Buffer once they have been paired with our indexing reagent and to store them for up to four weeks without degradation of data quality. This decouples the initial stages of sample collection and processing from downstream library preparation and so greatly increases flexibility in experimental design.

Finally, we demonstrated a unique property of our approach: the ability to selectively generate data from specific cell types within a heterogeneous mixture without using any kind of upstream cell type enrichment. This relies on the use of antibodies to drive pairing of our indexing reagent only to cells that carry a particular protein on their surfaces. This will enable experimental designs where only a given cell type is of interest from a mixed population without the need for complicated sample preparation nor wasting sequencing effort on unwanted cell types. Importantly, the targeted cell surface protein can be either endogenous (eg CD3E, CD19) or introduced experimentally to a subpopulation of cells (eg membrane-associated GFP). We believe that these selective capture methods will have great utility in basic research but also in translational applications such as quality control and patient monitoring for cell and gene therapy.

Taken together, these results demonstrate that Kinetic Confinement is a novel and effective approach to single cell RNA sequencing and that it is compatible with a range of sample types and experimental approaches. Furthermore, this simple solution-phase method will enable single cell genomics to be used with study designs that have, until now, required other less informative assays.

## Materials and Methods

### Cell line culture

HeLa (human epithelial cell line) and, NIH/3T3 (mouse fibroblast cell line) were separately cultured in T-75 flask with DMEM (4.5 g/L glucose, Gibco, ref. 11965092) supplemented with either 10% heat-inactivated foetal bovine serum (FBS, Gibco, ref. A5209402) or just bovine serum (BS, Gibco, ref. 26170-043) respectively, and 1% penicillin–streptomycin (ThermoFisher, ref. 15070063). Both cell lines were grown at 37 °C with 5% carbon dioxide (CO_2_) saturation and split when achieving a confluency of 80-90% for a maximum of 15 passages in total.

### Single cell preparation from adherent cell lines

Per cell type, at a confluency of approximately 80%, culture media was removed, and cells were washed twice with room-temperature (RT) DPBS (Gibco, ref. 14190144) for subsequent detachment using TrypLE™ (ThermoFisher, ref. 12605010) for 3 min at 37 °C inside the cell incubator. TrypLE™ was then inhibited by adding twice the volume of pre-warmed media; floating cells were collected in a fresh falcon tube and centrifuged at 200 x g for 3 min at RT. Supernatant was discarded whilst the pellet was resuspended in 1mL of cold DPBS + 0.04% BSA (Gibco, ref 15260-037), transferred to a 1.5 mL Eppendorf tube and centrifuged again at 200 x g for 3 min at 4 °C. Supernatant was once more discarded, pellet thoroughly resuspended in 1 mL of cold DPBS + 0.04% BSA and filtered through a 40 µm cell strainer (Flowmi™ ref. 136800040) into a new 1.5 mL tube.

### Preparation of human PBMCs from fresh blood samples

PBMCs were isolated by gradient centrifugation using Lymphoprep™ (StemCell, ref. 07851) according to manufacturer’s instructions. Briefly, freshly K3-EDTA-collected blood samples were mixed 1:1 with plain RPMI at RT, laid onto the recommended volume of Lymphoprep™, and centrifuged at 800 x g for 30 min at RT with acceleration/deacceleration switched off. The buffy coat was carefully transferred to a new 15 mL falcon tube and washed with RPMI at 400 x g for 5 min at RT. Subsequently, red blood cells (RBC) were lysed using RBC 10X Lysis solution (Invitrogen™, ref. 00-4300-54) according to the manufacturer’s protocol. The final PBMC pellet was washed twice with cold DPBS + 0.04% BSA at 200 x g for 3 min at 4 °C. Pellet was resuspended in cold DPBS + 0.04% BSA for viability and concentration assessment.

### Preparation of frozen whole-blood samples

To analyse 96 frozen whole blood samples in a single experiment (consisting of 2 samples from 48 individuals), samples were processed in batches of 24 samples from thawing to freezing the cell-CPair conjugate in the KCB reagent. All batches were processed in a single day with 45 minutes in between the start of each batch. Per batch, samples were processed as described below at IMU Biosciences (London).

Freshly lithium heparin-collected blood samples were chilled on ice and then thoroughly mixed in a 1:1 ratio with ice-cold freezing media (RPMI + 20% DMSO). Rapidly, up to 1mL of diluted blood was aliquoted per cryovial per donor, placed inside a cell freezing container CoolCell™ (Corning, ref. CLS432001) containing pre-cooled steel cores, stored in a -80 °C freezer until frozen and transferred into liquid nitrogen overnight.

Prior to the Kinetic Confinement assay, cryovials were rapidly thawed on a 37 °C heat block. Thawed samples were individually transferred to new 15mL falcon tubes per donor, completed to 10 mL with pre-warmed RPMI and centrifuged at 500 x g for 5 min at RT. Cell pellets were resuspended and red blood cells were lysed using BD Pharm Lyse buffer (BD Biosciences, ref 555899) for 15 minutes. For scRNAseq, an aliquot of cells per sample was centrifuged at 500 x g for 5 min at RT and used for dead cell removal (Miltenyi Biotec, ref. 130-090-101) using the autoMACS® Pro Separator following manufacturer’s instructions. From this point onwards, all steps were performed on ice with pre-cooled buffers. Negatively eluted viable cells were then washed twice with cold DPBS and centrifuged at 500 x g for 5 min at 4°C before assessing both viability and cell count using the Luna–FX7™ cell counter (Logos Biosystems). Samples were appropriately diluted in DPBS + 0.04% BSA. Once all 96 samples had been transferred to KCB and frozen, all batches were processed together to sequencer-ready libraries as a single set of samples in 96-well plates. In parallel to the preparation for scRNAseq, the remaining cells were stained for flow cytometry as described below.

### Flow cytometry staining and acquisition

Cells were resuspended in a solution of viability dye, then incubated for 15 min at room temperature to label dead cells. Cells were then washed and labelled with a panel of fluorescently-labelled antibodies targeting cell surface markers. First, cells were resuspended in a mixture of chemokine receptor-specific antibodies and incubated at 37 °C for 15 minutes. Second, a mixture of the remaining antibodies was added to the cells and they were incubated for 45 minutes at room temperature. Third, cells were fixed with Lysing Solution (BD, ref 349202) before being washed twice and resuspended in FACS buffer. Stained cells were acquired with an ID7000 full spectrum flow cytometer (Sony). QC check was performed daily in advance of sample acquisition using AlignCheck particles and 8 peak beads (Sony, ref AE700510 and AE700522). Full spectrum signals were unmixed and fcs files were exported to Amazon S3.

### Single cell preparation for mouse splenocytes

Fresh mouse spleen samples were purchased from Charles River Laboratories and shipped in Tissue Storage Solution (MACS™ ref. 130-100-008) at 4 °C. Upon arrival spleen samples were processed individually; each spleen was separately rinsed with cold DPBS, cleaned from connective tissue, and placed inside a 40 µm cell strainer (Corning, ref. 431750). The strainer containing the spleen was placed on a new petri dish filled with 10 mL of ice-cold RPMI. Using the plunger of a syringe, the spleen was gently macerated against the strainer until fully dissociated. Leftover connective tissue was discarded and splenocytes collected in a fresh 15mL falcon tube for centrifugation at 500 x g for 5 min at 4 °C. Subsequently, two washes with ice-cold DPBS + 0.04% BSA were performed before assessing viability and cell concentration.

### Viability and cell concentration assessment

Cell concentration and viability were measured using a combination of AO/PI staining using the CellDrop instrument (DeNovix). Samples with < 90% viability underwent Dead Cell removal (MACS™, ref. 130-090-101) following manufacturer’s instructions. Cell suspensions were diluted with cold DPBS + 0.04% BSA to achieve a homogeneous concentration of 425 cells/µl of PBMCs or 250 cell/µl per cell line with >90% viability. For cell lines, individually diluted HeLa and NIH/3T3 cell suspensions were mixed at 1:1 ratio for a combined cell concentration of 250 cells/µl . The final concentration and viability of cell suspensions were confirmed with AO/PI in the CellDrop before being used in the SimpleCell™ assay.

### Single cell RNA sequence library construction using Kinetic Confinement

Single cell suspensions were processed into single cell RNA-seq libraries using the SimpleCell™ 3’ Gene Expression Assay (CS Genetics, ref 1000393) as per manufacturer’s instructions. 5000 NIH/3T3 and HeLa cells in a 1:1 ratio or 8500 PBMCs were paired with CPair and frozen on dry ice in Kinetic Confinement Buffer. Frozen samples were kept at -80 °C for a minimum of 40 minutes until library preparation. The MiniAmp Plus (Applied Biosystems) was used for cell lysis and mRNA indexing (95 °C for 15 sec, 55 °C to 40 °C decreasing at 1°C /6 sec, 35 °C to 20 °C decreasing 1 °C /9 sec, 20 °C hold), Reverse Transcription (20 °C for 2 min, 30 °C for 2 min, 40 °C for 2 min, 50 °C for 15 min, 55 °C for 15 min, 98 °C for 3 min, 4 °C hold), RNaseH treatment (37 °C for 20 min, 65°C for 5 min, 4°C hold), Second Strand Synthesis (37°C for 20 min, 65°C for 5 min, 4°C hold) and Proteinase K treatment (37°C for 15 min, 55°C for 10 min, 4°C hold). Mag-Bind TotalPure (Omega Bio-tek, ref. M1378-00) were used to clean up the double-stranded cDNA. cDNA was amplified with 12 cycles (cell lines) or 13 cycles (PBMCs) using the MiniAmp Plus (98°C for 45 sec, cycled 13 or 12x : 98°C for 15 sec; 60°C for 30 sec; 72°C for 30 sec, 72°C for 1 min, 4°C hold). Amplified DNA was size selected using a double-sided DNA cleanup (upper 0.5x and lower 0.7x) using Mag-Bind TotalPure beads. The library was enriched according manufacturer’s instructions and Illumina UDIs were added via an indexing PCR with 6 cycles (cell lines) or 7 cycles (PBMCs) using the MiniAmp Plus (98°C for 45 sec, cycled 6 or 7x: 98°C for 15 sec; 60°C for 30 sec; 72°C for 30 sec, 72°C for 1 min, 4°C hold). Indexed libraries were cleaned up using Mag-Bind TotalPure beads. Library size was analysed using the Tapestation 4200 (Agilent, ref G2991BA) with the HS-NGS High Sensitivity 474 kit (Agilent, ref DNF-474-0500). Libraries were quantified using qPCR with the NEBNext® Library Quant Kit for Illumina (NEB, ref E7630L). Pooled libraries containing 2% PhiX were loaded on a NextSeq2000 (Illumina) at 1.3 nM and sequenced with the following read lengths: Read 1 - 92 cycles, Index 1 - 10 cycles, Index 2 - 10 cycles, Read 2 - 26 cycles.

### Computational processing of sequencing data

The CS Genetics scRNAseq data processing pipeline operates on paired-end sequencing data. Firstly, the 13 bp CPair barcode is identified in Read 2 and matched against a list of known, allowed barcode sequences. Barcodes with 1 Hamming distance are corrected to the closest known sequence whilst those with a greater Hamming distance are discarded. Retained read pairs are processed to remove non-transcriptomic sequences introduced during library construction. Additionally, homopolymeric sequences longer than 14 nt are removed from the 3′ end of Read 1. Finally, any polyA sequences greater than 14bp or polyA sequences greater than 12bp followed by ‘CG’ are removed (including all sequence 3’ of that sequence). Read pairs with R1 longer than 4 bp after trimming are retained.

Processed reads are mapped to a genomic reference using STAR^14^ and filtered to permit a maximum of 3 mismatches to the reference. Reads are associated to genes using featureCounts^15^. Reads are separated into uniquely mapped reads (UMRs) with a single alignment, and multimapping reads with more than one equally likely alignment. UMRs are first annotated against ‘transcript’ features within the reference GTF. Reads categorized as ‘Assigned’ are directly associated with the transcript’s parent gene. Reads categorized as ‘Unassigned_Ambiguity’ are used as input to featureCounts a second time and annotated against ‘exon’ features within the GTF. Again, ‘Assigned’ genes are associated with the exon’s parent gene. All other reads are discarded. For multimapping reads, if all possible alignments for a given read are ‘Assigned’ to a single gene, that read is directly associated with that gene. If a read is ‘Assigned’ to multiple different genes, or if there are ‘Unassigned_Ambiguity’ alignments, the alignments are passed onto the exon annotation with the same logic applied to assign reads to genes.

Reads tagged with the same cell barcode and mapping to the same genomic start coordinate, are considered PCR-duplicates and deduplicated using UMI-tools (Smith et al. 2017). Deduplicated reads that are assigned to a gene annotation are considered as counts. For single species datasets, CPair barcodes with counts above a per-sample, dynamically determined threshold are defined as representing cells. To determine this dynamic threshold, a KDE smoothing is applied to a histogram of the log10-transformed per-barcode counts. The count threshold is then defined as the first minimum of the KDE above 10^2^. For mixed-species datasets two thresholds are determined. For each species, the same KDE smoothing method is used, but applied only to those CPair barcodes that have a majority of counts for the species in question, and only using counts associated to genes from that species. Single species CPair barcodes are then defined as those barcodes with counts above one threshold and below the other. Thus, a CPair barcode is identified as the first species if it has counts above the first species threshold but below the second. Barcodes with counts below both thresholds are classified as noise. Barcodes with counts above or equal to both thresholds are considered multiplets. Gene expression is estimated for a cell-associated CPair barcode by counting the number of deduplicated reads assigned to each gene for that CPair barcode.

### Computational analysis of gene expression data

Count matrices generated from the raw sequencing data were used as input into Seurat^16^ or Scanpy^17^ for analysis. Preprocessing and QC steps were performed to remove multiplets and poor-quality cells based on dynamic thresholds for total counts, genes and mitochondrial read percentage. Typically, cells within 95% of the respective distributions were kept for subsequent processing. Log-normalisation followed by data scaling was applied, while controlling for differences in mitochondrial read percentage.

Dimensionality reduction for plotting was performed using Uniform Manifold Approximation and Projection (UMAP)^18^ while cell clusters were identified using the graph-based Louvain algorithm. Cell type identities were annotated using differentially expressed marker genes that were associated with each cluster along with knowledge of expected subtype marker genes.

## Supplemental material

**Supplementary Figure 1.**
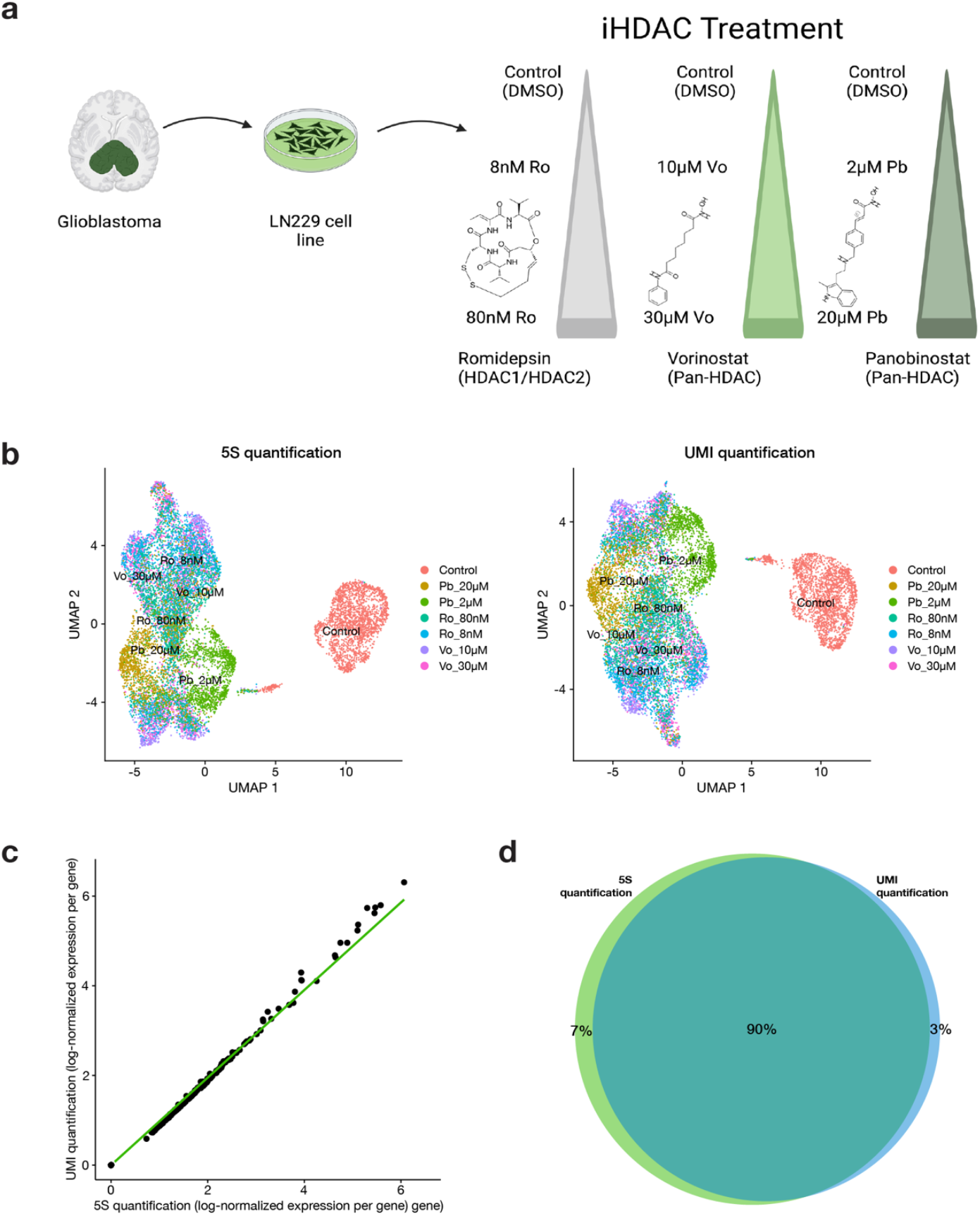
Comparison of gene expression quantification using Second Strand Synthesis Start Sites (5S) and unique molecular identifiers (UMIs). **a**, Experimental design. Glioblastoma cell line LN229 cultures were treated with varying concentrations of three different histone deacetylase (HDAC) inhibitors and then analysed using Kinetic Confinement-based single cell RNAseq. **b**, Comparison of UMAP dimensionality reduction and graph-based clustering results using gene expression matrices generated by each quantification method. Very similar population structures are observed in both cases. **c**, Correlation of log-normalised gene expression values for each method. Correlation coefficient: >0.99, slope ∼1.0. **d**, Comparison of between-cluster differentially-expressed genes (DEGs) identified using each gene expression quantification method. 5S quantification revealed 3650 DEGs in total while UMI quantification revealed 3437.

**Supplementary Figure 2.**
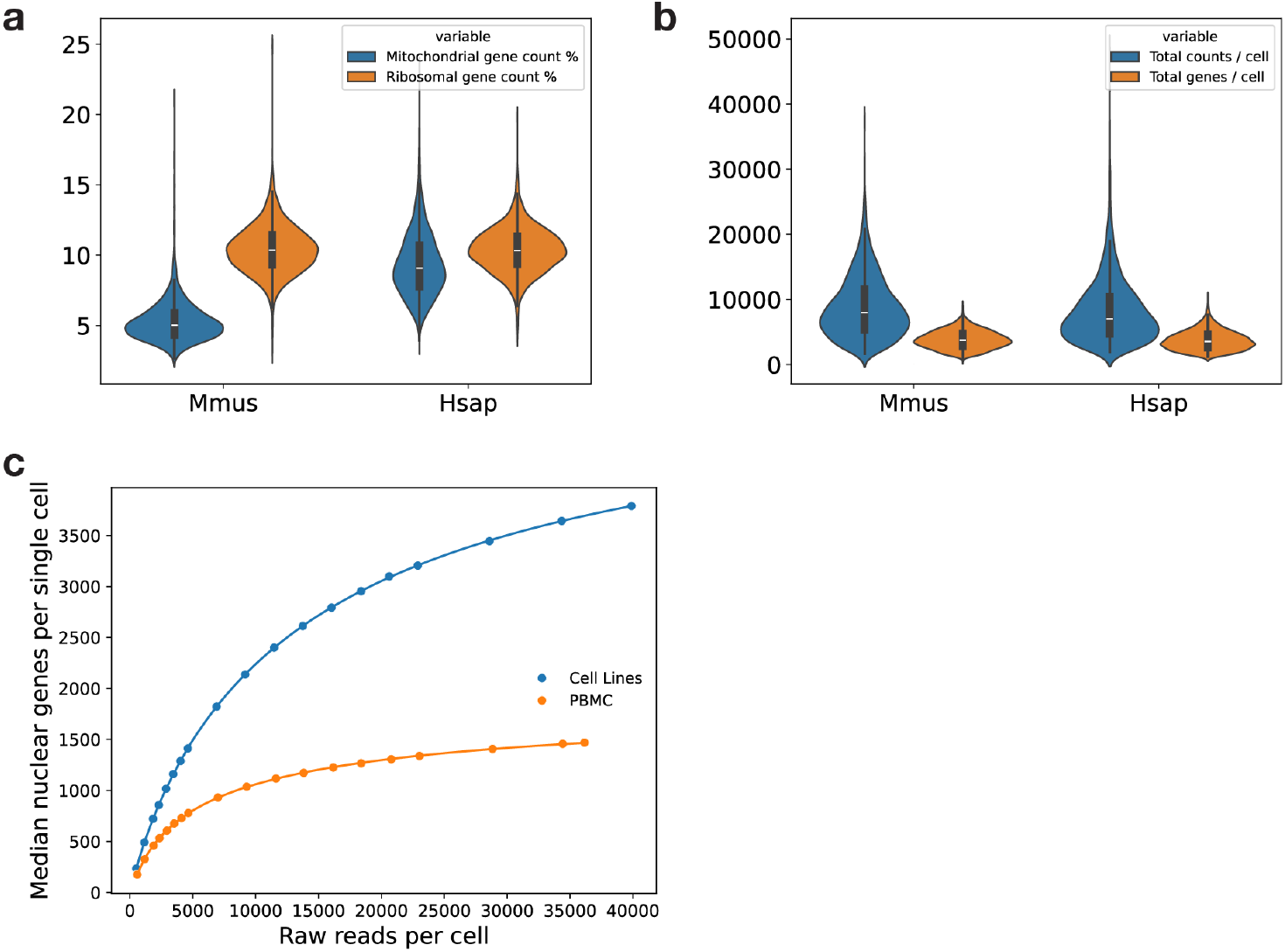
Performance metrics from mixed-species experiments as, Per-species distributions of per-cell quality control metrics: percentage of reads derived from the mitochondrial genome (blue), and percentage of reads derived from genes encoding ribosomal proteins (orange). **b**, Per-species distributions of the number of per-cell 5S counts (blue) and genes detected (orange). **c**, Impact of downsampled read depth on the number of genes detected in each single cell for mixed-species cell lines (blue) and PBMC (orange) samples.

**Supplementary Figure 3.**
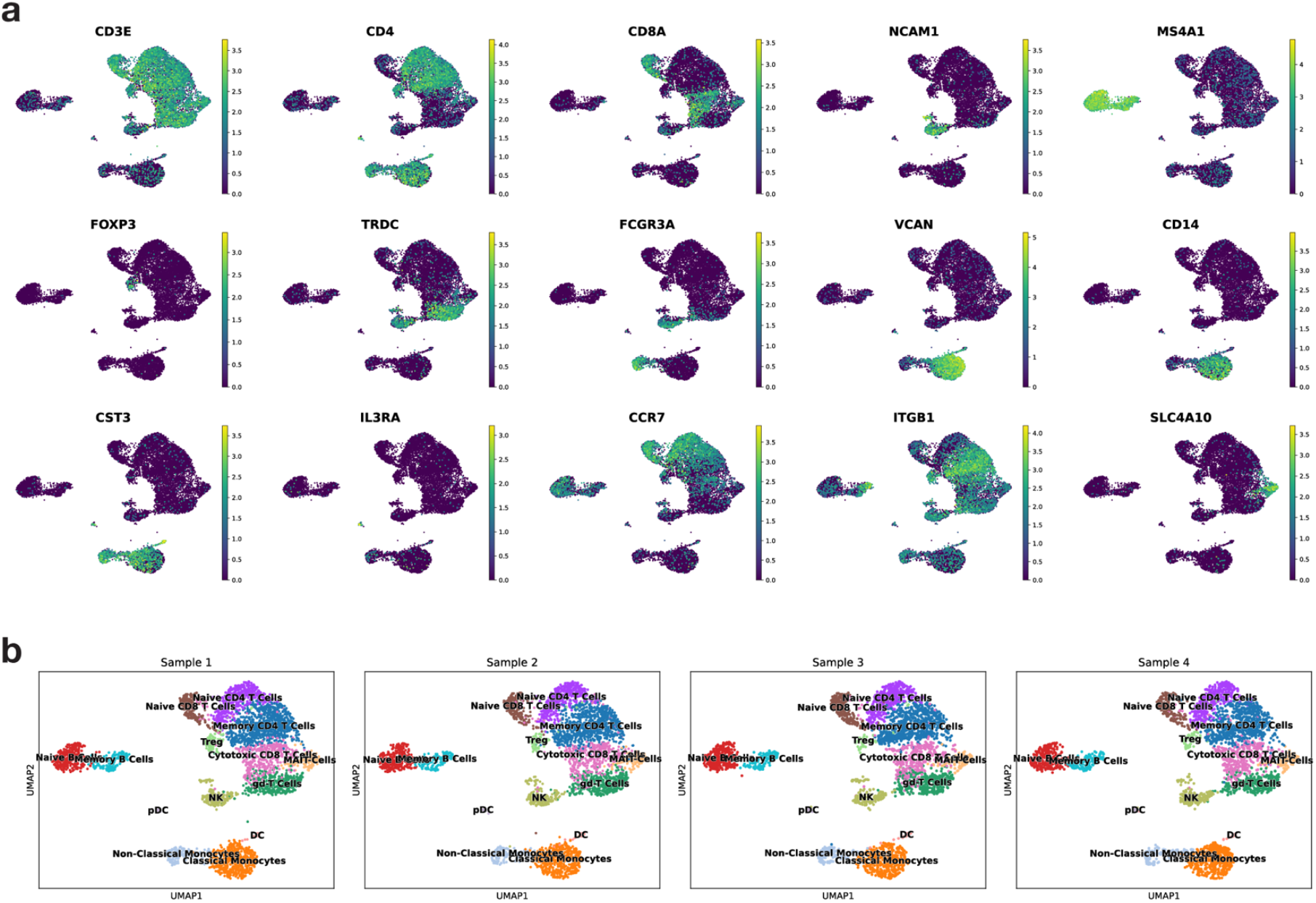
Detection of cell subtypes within heterogeneous mixtures from blood a, Expression of selected marker genes within UMAP plots generated from PBMC samples **b**, UMAP plots for each of four PBMC samples that were combined for computational analysis

